# Testing for pre- and post-copulatory inbreeding avoidance in the flour beetle *Tribolium castaneum*

**DOI:** 10.1101/2025.10.10.681665

**Authors:** Fathi Ali Attia, Tom Tregenza, Ramakrishnan Vasudeva

## Abstract

Inbreeding depression poses a significant threat to fitness in many taxonomic groups, particularly those where close relatives are likely to come into contact. Studies have suggested that females may mate with multiple males to reduce inbreeding through post-copulatory processes of closely related males. In our mating trials, we examine potential pre- and post-copulatory inbreeding avoidance in the red flour beetle *Tribolium castaneum* using behavioural observations and subsequently, through the use of phenotypic mutant markers to assign parentage. As expected, mating rates were largely limited by females. Contrary to expectations, males courted and mated with related females more frequently than with unrelated females. Unrelated males sired a greater proportion of the offspring of doubly mated females, but this was explained by the higher survival of their offspring to adulthood. When the lower survival of inbred offspring was accounted for, there was no difference in the estimated fertilisation success of related and unrelated females. We show that last male sperm precedence declined over 5 weeks, suggesting that the cause of sperm precedence was primarily displacement of previously stored sperm, with evidence for a limited degree of sperm stratification within the spermatheca.

## 1. Introduction

Inbreeding depression, the loss of fitness resulting from homozygosity, can occur within otherwise outbred populations as a result of matings between relatives. It is caused by the expression of rare deleterious recessive alleles and loss of heterozygosity, although the relative contributions of these factors are not well understood [1]. Inbreeding depression in captive animals is well documented [2], and there are a growing number of examples from natural populations [3]. Many vertebrates actively disperse from their natal sites, and manipulative studies suggest that this behaviour may be driven by benefits of reduced inbreeding [3]. Dispersal can relax selection for active inbreeding avoidance behaviour [4]). In species with dispersal strategies, the risk of inbreeding may be too low to drive the evolution of sibling recognition mechanisms [5]. In some species, individuals not only distinguish between kin and non-kin, but may even discriminate among close kin and distant kin [reviewed by 6]. Grosberg and Hart [7] reported that an allorecognition polymorphic gene may regulate the reproductive interaction between mates or between gametes to avoid inbreeding. Overall, the available evidence suggests that costs of inbreeding represent a significant selection pressure across a broad range of taxa.

Inbreeding avoidance has been proposed as a potential explanation for the prevalence of polyandry in groups where matings do not provide direct benefits to females [8, 9] and inbreeding costs are high [10]. Although no evidence for such a process was found in shrews, for which the hypothesis was originally proposed [11], evidence from 3 species of shorebirds [12] and from blue tits (*Parus caeruleus*) [13] indicates that females utilise extra-pair copulations to reduce inbreeding. Furthermore, studies of guppies and 2 species of field crickets indicate that multiply mating females can avoid costs of inbreeding through increased fertilisation success of unrelated males [14-16], suggesting it would be worth looking for similar effects in other species.

Studies of potential precopulatory and postcopulatory inbreeding avoidance in *Drosophila melanogaster* have failed to find any evidence for it [17, 18]. Indeed, evidence for inbreeding avoidance in invertebrates is limited, but this may also reflect the fact that it has rarely been studied [3]. It is most likely to be found in species whose ecology leads to high risks of encountering closely related individuals. Such species will include those with strong population structure and philopatry, and also those that are frequently exposed to founder events. Stored product pests represent such a group since they exist on resources that are often ephemeral and rely on their ability to locate and colonise new food patches. Such colonisation events may frequently involve a small number of individuals arriving at a new patch, with concomitant risks of mating amongst relatives.

The red flour beetle, *Tribolium castaneum* is a stored product pest found in flour stores throughout the temperate and tropical zone. It has been used as an experimental model system across many fields of ecology and evolution [19]. Populations have been shown to experience inbreeding depression, suffering reduced egg laying rate and hatchability [20], sperm length, testes volume, reproductive output and lifespan, with variation between populations in the extent of inbreeding depression [21]. In *T. castaneum*, it has been shown that inbreeding increased levels of female promiscuity compared with noninbred cohorts [22]. *T. castaneum* is known to exercise mate choice using scent cues to distinguish between mates on the basis of female maturity and previous mating history [23, 24], and more attractive males are known to have higher P_2_ [25], so there is potential for olfactory kin recognition. The fertilisation success of male *T. castaneum* was not depressed under strong inbreeding, but inbreeding affected the outcome of sperm competitiveness [26]. Whether copulations result in complete failure of fertilization was not influenced by the relatedness of mating pairs [27].

In this study we examine costs of inbreeding in *T. castaneum* and investigate whether adults can distinguish between related and unrelated potential mates, whether they discriminate against relatives when choosing to mate and whether females can bias paternity towards unrelated mates when multiply mated using two different strains.

## 2. Methods

Experimental beetles were sexed and isolated as pupae into single-sex groups of a standard age [28]. Pupae were kept individually in a 2cm x 2cm cell of a 5x5 cell plastic box with a mixture of 95% organic flour and 5% dried yeast in a dark incubator at 30°C and approximately 67% relative humidity (standard rearing conditions). Pupal isolation was crucial as males can engage in same sex behaviours and become sperm limited [29]. The eclosion date of each beetle was recorded to minimise age effects during mating trials (see, online supplementary material). All beetles used in this experiment were between 9 and 21 days post-eclosion, and the maximum age difference between any beetles in a trial was 2 days. Two strains were used, the wild type GA1 strain (Georgia 1, with filiform, wild type antenna) and the reindeer strain (Rd strain, with swollen antenna, see [30], which carries a recessive dominant marker that causes the antennae to have a pronounced terminally clubbed type, reduced and easily distinguished phenotype from the wild type antenna [30]. Both strain of beetles were provided by Dr R. Beeman, US Grain Marketing and Production Research Centre and maintained in the laboratory under identical standard rearing conditions.

### 2.1 Test for pre-mating inbreeding avoidance (behavioural)

Interactions between GA1 strain beetles were observed in an arena that was a 2cm x 2cm cell. The bottom of the cell was covered with filter paper to provide traction for the beetles and care was taken so that the beetles were unable to climb the walls. The small interacting arena allowed frequent contacts between individuals that could be observed. In this arena, we recorded the focal male’s interactions with a beetle of the same sex or the opposite sex, including whether the focal male attempted to climb onto the other individual (mounting attempt). During this phase, no antagonistic behaviour was observed and resistance to mounting consisted simply of attempts to walk away from the mounting male. Full sibling (‘sib’) families were reared with offspring separated, sexed at the pupal stage and kept individually. We examined two situations: a) male mate choice, in which two unrelated (‘non-sib’) females were placed with a single male who was the full sibling of one of them (27 independent trials); and b) female mate choice in which two unrelated males were placed with a single female who was the full sibling of one of them (30 independent trials). All individuals were virgin at the beginning of the trial. Both individuals of the same sex were marked with different colours of permanent marker pen (colours allocated at random) to allow them to be identified. Marking is known not to interfere with reproductive output of interactions that lead to copulations [31].

#### Male mate choice traits

We quantified contacts initiated by focal males (counts), number of mating attempts towards sib versus non-sib females and mating duration (seconds) within these assays

#### Female mate choice traits

Behavioural observations allowed us to quantify contacts initiated and mating duration towards sib versus non-sib individuals.

Pre-copulatory behavioural observations allowed us to record the number of mating attempts between sibs and non-sibs of the opposite sex that took place over a 30-minute period. Additionally, we recorded the duration of the first mating. Our observations revealed that females only occasionally initiated contacts with males, therefore in both studies we defined mating attempts as occasions when the male inspected the female’s head or abdomen with his antennae or maxillary palps and attempted to dorsally mount the female [23]. Attempts that were followed by mating were considered as an attempt and a mating. Mating durations of less than 40 seconds were considered as attempts since such short copulations rarely result in successful fertilisation [32].

### Post-mating inbreeding avoidance

We used a similar design to that used to investigate post-mating inbreeding avoidance in field crickets by Tregenza and Wedell [16]. However, in this study we compared fertilisation success of GA1 males that were full siblings (sib) with a GA1 female with the fertilisation success of unrelated (non-sib) Rd males. We used these two strains as previous studies have found no difference between them in their reproductive output [33], also see [30]. A preliminary study (data from [33]) of the number of offspring produced by ten GA1 females mated with either a sib or non-sib GA1 male, or an Rd strain male revealed that sib crosses produced significantly fewer offspring than crosses involving Rd males (estimate: -0.39, z=-3.4, P<0.0001, Fig. 2 panel D). Non-sib GA1 male and Rd male crosses with GA1 females produced similar numbers of offspring (estimate: -0.16, z = -1.25, P = 0.209, Fig. 2 panel D). Our design has the limitation that it confounds non-sib with ‘non-strain’; ie. the non-sib crosses are an Rd male and a GA1 female whereas the sib crosses are a GA1 male and a GA1 female. This means that if we find a difference between the post-mating reproductive success of sibs and non-sibs, we need to be aware that this difference could arise if there are interactions between strains that affect fertilisation probably or offspring viability (even though the available evidence suggests that such differences do not exist).

Twenty-five full-sib families were reared and used in forty-two experimental blocks, with no family used in more than 2 blocks. Each block consisted of four GA1 females each mated to two males in one of the following combinations: either to two full-sib GA1 males (SS); two non-sib Rd males (NN); a sib GA1 male followed by a non-sib Rd male (SN); or to a non-sib Rd male followed by a sib GA1 male (NS). Within a block, all ‘GA1’ and ‘Rd’ males were full-sibs and of similar age groups.

In each mating trial, the male and female were placed together in the same chamber as described above for the pre-mating study and observed until a mating lasting more than 40 seconds took place. As soon as the pair separated, the male was removed and the female was left individually in the cell for 24 hours. Females are reluctant to lay eggs in the absence of a suitable substrate, and we did not observe any egg laying over this period. After 24 hours the second male was placed with the female, and the pair were observed until they mated for at least 40s. All females were virgin at the start of the experiment and mated with both males. All males were used only once and were mated once to a stock female on the day before they were used in the experiment. When the female’s second mating finished, she was kept in a 100ml pot containing 40ml of flour in a dark incubator at 30°C and 67% RH. After one week, the female was removed and the pot returned to the incubator under the same climate regime as above. The number of adult offspring were counted 45 days after the last day of oviposition, which was before any eggs laid by the first emerging offspring could themselves reach adulthood. Additionally, in 17 randomly chosen blocks, females were kept for 5 further weeks. Their offspring were collected and counted each week by moving the female to a new pot every week for 5 weeks to calculate P_1_ (offspring fathered by the first male) and P_2_ (offspring fathered by second the male). This design allowed us to examine any changes over time in P_1_ and P_2_.

## Statistical methods

All analyses were carried out R (v. 4.4.1) [34] in R Studio (2024.09.0-375) [35]. Plots were produced using “ggplot2” package v4.0.0 [36]. Data summary was obtained by using the “RMisc” package v1.6 [37]. Model diagnostics were performed using ‘performance’ v0.15.1 [38] and ‘see’ v0.12.0 packages [39].

Our preliminary study using GA1 beetles to study offspring productivity between ‘Rd’ and ‘sib’ ‘non-sib’ male:female crosses was analysed using a negative binomial generalized binomial model using the MASS package v7.3-65 [40] with offspring counts as the response and ‘treatment ID’ as the fixed factor.

Pre-copulatory mate choice behaviours in both sexes were analysed using glmmTMB [41] with contacts initiated (counts) by male, female and mating duration in seconds (log transformed) towards sib versus non-sib individuals as response variable and with appropriate error distributions. Count data were fit with a ‘poisson’ or ‘negative binomial’. Mate type (‘sib’, ‘non-sib’) as explanatory variables and replicate ID as random factor.

## Results

### Pre-mating inbreeding avoidance

Our 27 male mate choice trials resulted in an average of 37 ± 0.46 (mean ± S.E.) mating attempts and 4.90 ± 0.31 matings over the half hour period. A heterogeneity χ^2^ test of the trials showed no significant differences between replicates in the numbers of attempts and numbers of matings (χ^2^ = 21.1, *P* = 0.74 and χ^2^ = 29.9, *P* = 0.27, respectively). We found that when males were given a choice, they attempted to mate with their siblings (589 times), significantly more frequently than with non-sibling females (411 times), (estimate: 0.35, z = 5.59, P<0.0001, Fig. 1 panel A). The overall difference between the number of mating attempts by males with their sibling (77 times) and with a non-sibling (55 times) was significant with a greater number of mating attempts directed towards female sibs (estimate: 1.00, z = 2.76, P = 0.005, Fig. 1 panel B). Mating duration of sibling and non-sibling males revealed no statistical difference (estimate: 0.32, z = 0.81, P = 0.42, Fig. 1 panel C)

**Fig. 1.**
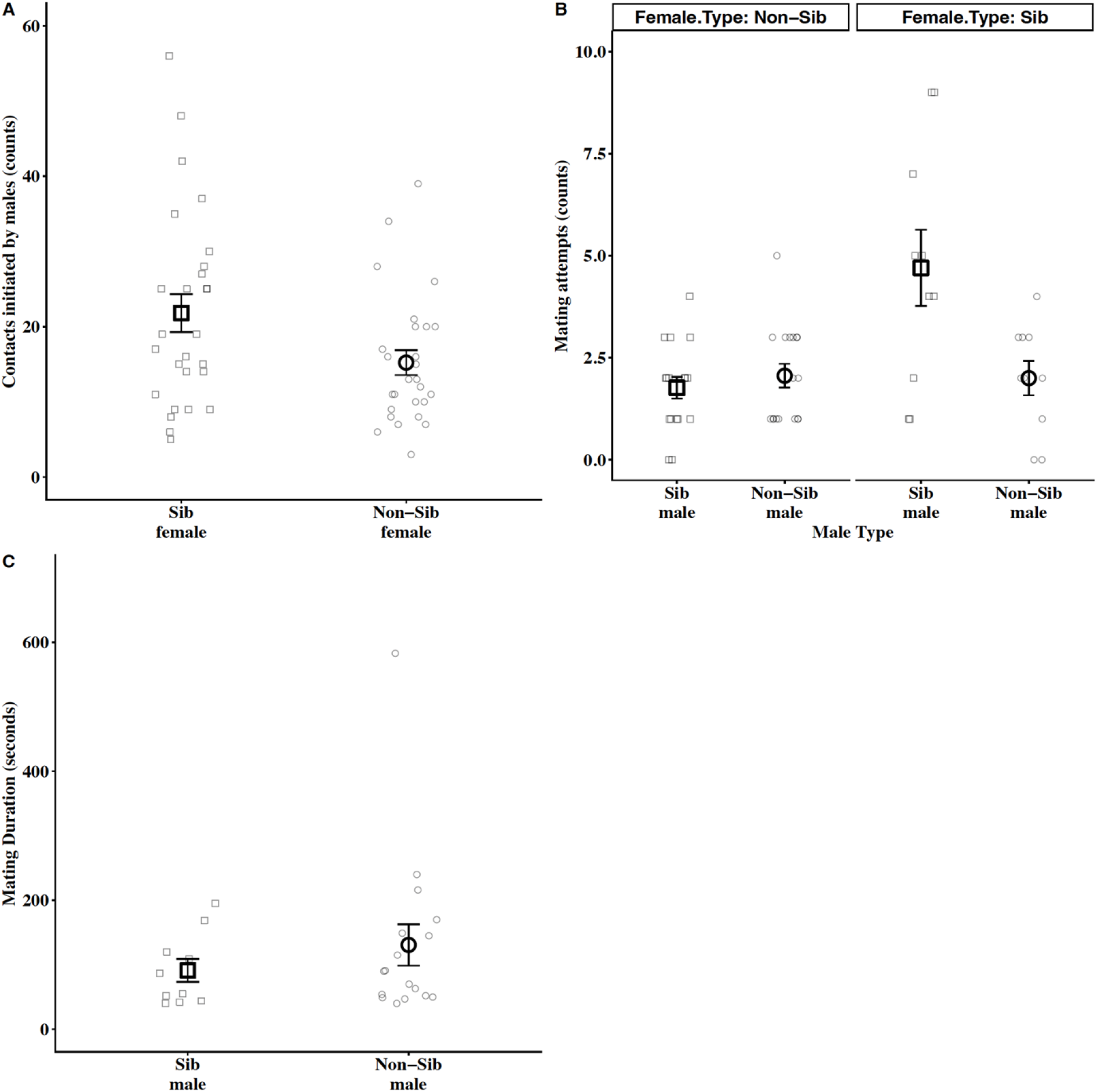
Pre-mating male behavioural traits when interacting with either a full-sibling or a non-sib female. Panels A–C represent data (mean ± SE, full-siblings [square] and non-sibs [circle]). Raw data are denoted as jittered points. In panel A, contacts initiated by the males towards full-sib or non-sib female was measured. Panel B measures mating attempts by males towards either a full-sib or non-sib female. Mating duration (panel C) when the male is a full-sib versus non-sib was measured.

In our female choice trials, observations revealed that females only occasionally initiated contacts with males (mean of 2.5 contacts per 30 minute observation period; estimate: 0.21, z = 0.91,P = 0.36, Fig. 2 panel A). The two males in each trial together attempted to mate with the female 17.7 ± 0.51 times, with 3.81 ± 0.25 matings taking place over the 30 minutes. Additionally, males were observed to engage in homosexual pseudo-copulations with other males with a similar frequency to their copulations with females (mean of 3.74 ± 0.23 per half hour). Data on the number of contacts between males of each type and the female, revealed significant heterogeneity between trials (heterogeneity χ^2^ = 148, d.f. = 29, P<0.001). The number of contacts between sibling and non-sib males and the female overall were very similar (257 and 263 respectively; estimate: 0.08, z = 0.43, P = 0.66, Fig. 2 panel B). The number of matings between females and sibling and non-sib males (55 and 57 respectively) were pooled (heterogeneity χ^2^ = 39.7, d.f. = 29, *P* = 0.11) revealed no significant difference between the two (χ^2^ = 0.03, d.f. = 29, *P* = 0.85).

**Fig. 2.**
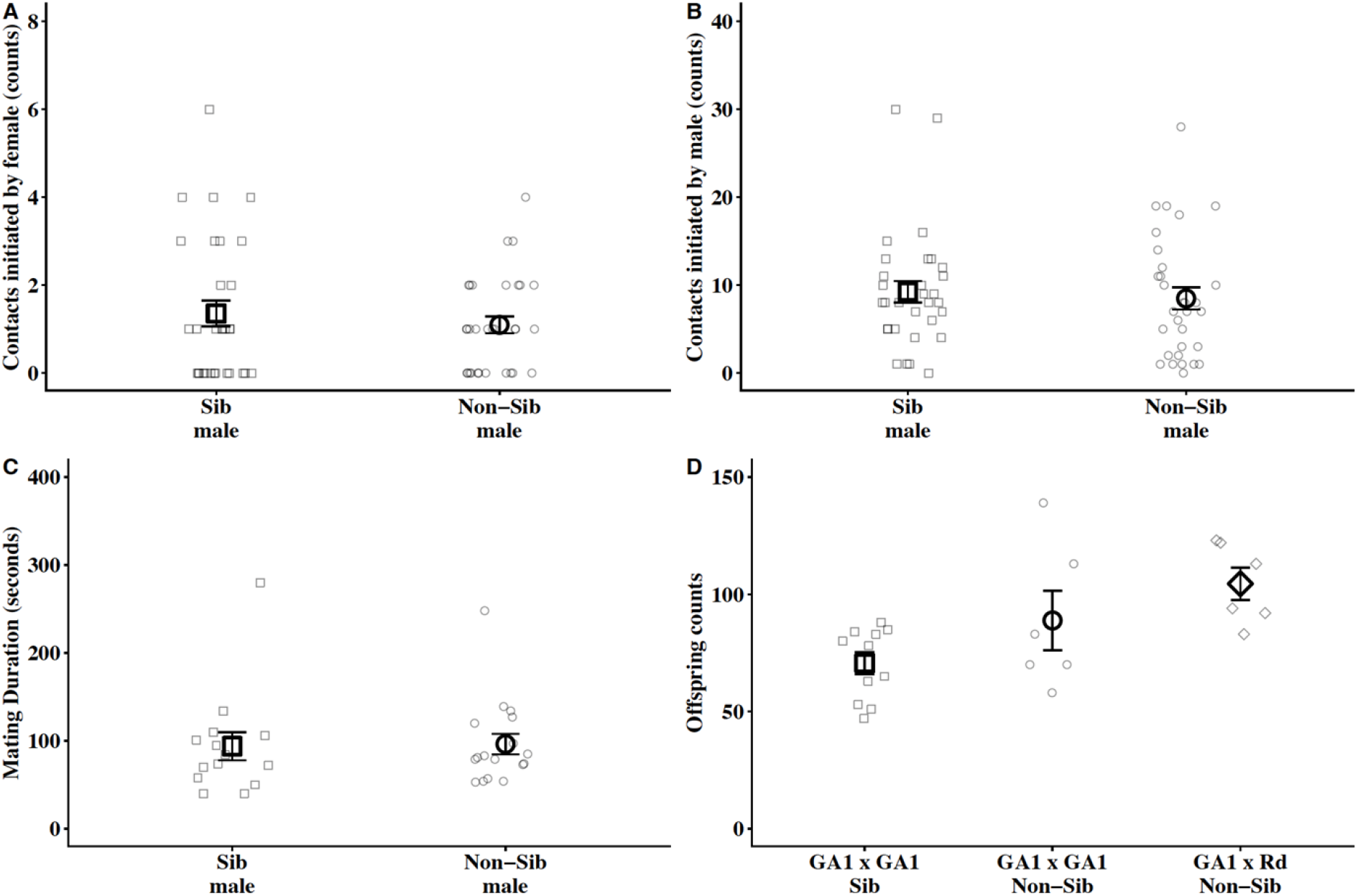
Pre-mating female behavioural traits (panels A–C) when interacting with males that were either a full-sibling or a non-sib to the focal female. Panels A–D represent data (mean ± SE) with a filled large central point indicating treatment average (shapes indicate full-siblings [square] and non-sibs [circle, diamond]). Jitters, fainter scattered points, around the central point indicate the raw data. In panel A, contacts initiated by the focal female towards full-sib or non-sib male was measured. Panel B denotes contacts initiated by males (counts) within female choice trials. Panel C indicates mating duration and panel D quantifies offspring produced by full-sibling GA1 pairs versus when paired with an unrelated GA1 and Rd non-sib males.

Across both experiments, first matings had a mean duration of 104 seconds. This is short compared to the observation period of 3600 seconds, so the number of matings will not have a large effect on the number of mating attempts that can be made. There were no differences in mating duration between sibling and non-sibling matings (estimate: 0.10, z = 0.45, P = 0.64, Fig. 2 panel C). Comparing numbers of attempted and actual heterosexual matings between treatments, reveals that number of contacts is lower when there are two males and one female (log transformed data, *t* = 5.15, d.f. = 56, *P*<0.001) but that numbers of matings are not significantly affected by the sex ratio (square root transformed data, *t* = 1.70, d.f. = 56, *P* = 0.094), hence the proportion of contacts that resulted in a mating differed between the sex ratio treatments (1 male, 2 females: 1/8.9 attempts successful, 2 males, 1 female: 1/5.9 attempts successful; log transformed data, *t* = 2.82, d.f. = 56, *P* = 0.007).

### Post-mating inbreeding avoidance

According to our earlier study [33], there fewer adult offspring were produced by GA1 females mated to GA1 full-sibling males compared to GA1 females mated to Rd non-sibling males (mean ± S.E.) number of offspring = 70.64 ± 4.6 and 104.5 ± 6.9 for GA1 sibling and Rd males respectively *z* = -3.4, *P*<0.0001, Fig. 1 panel D). This indicates that GA1 females do suffer costs of mating (lower offspring viability when the mate with related males as opposed to unrelated, non-strain Rd males.

The difference in fecundity of females in the 4 treatments (SS, NN, NS and SN) is shown in figure 3, panel A. These data include females that did not receive any sperm despite their two copulations and produced no offspring (10, 4, 4 and 3 females for SS, NN, NS and SN treatments respectively, χ^2^ = 5.9, P = 0.12), and females that received sperm from one male but not the other (which cannot be distinguished from double fertilised females in the NN and SS treatments). Failure to transfer sperm at mating is common and its frequency does not differ substantially between within and among strain crosses [32]. The difference in the offspring produced over one week was significant between sib-sib (SS) and non-sib (NN) pairs (estimate: -0.50, z=-2.35, P<0.01, Fig. 3 panel A).

**Figure 3.**
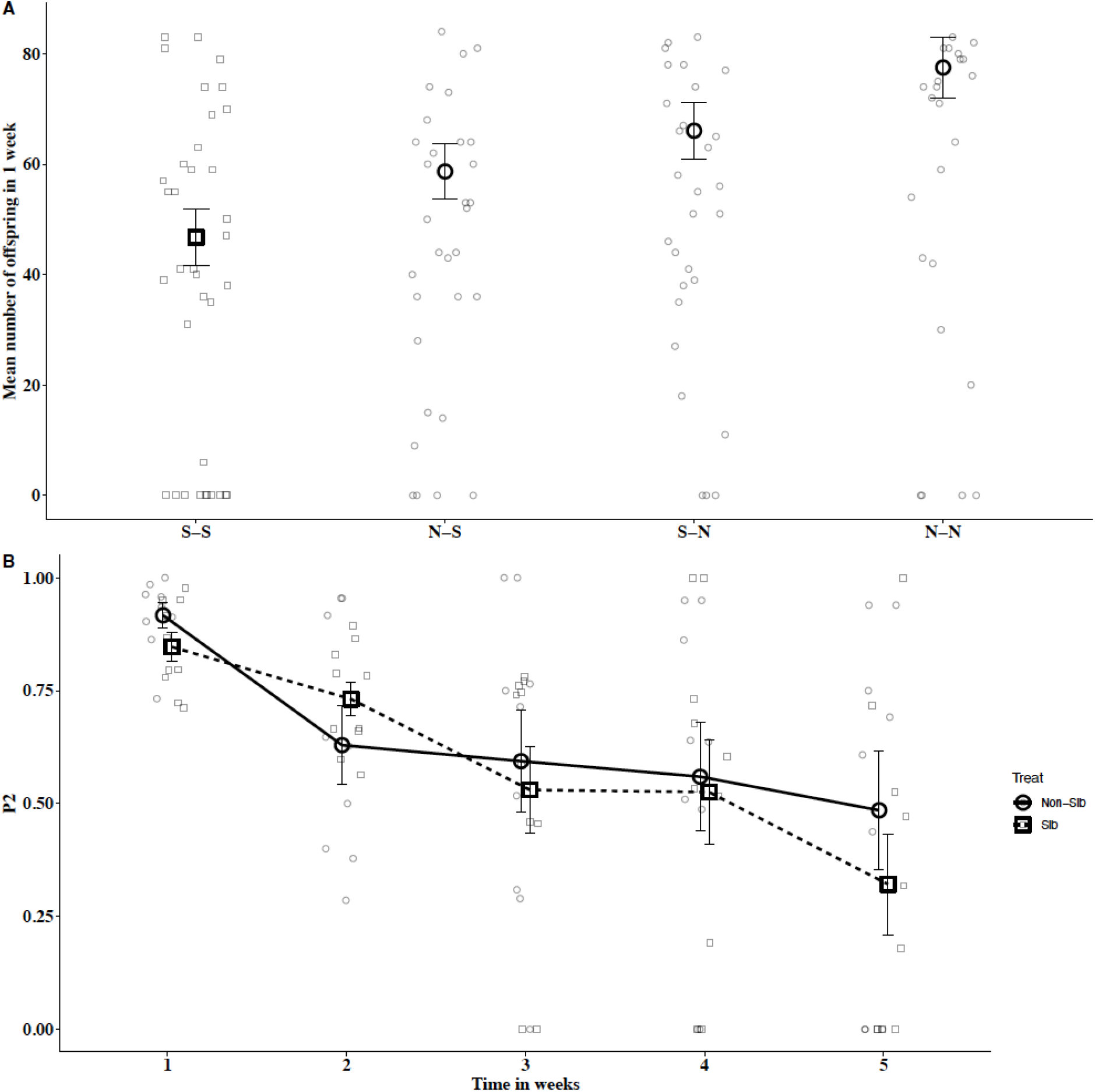
Panel (A) The mean number of adult offspring produced by females mating to differing combinations of related and unrelated males, *n* = 42 blocks of 4 sibling females, one per treatment. Panel (B). P_2_ ± S.E. of sibling and non-sibling males over 5 weeks adjusted to take into account the lower survival of offspring from sibling males (see text). Figure illustrates the decline in P_2_ over time.

The total number of offspring sired by the first and second male to mate with a female was calculated for those females that produced offspring from both males and ceased offspring production during the 5 week period. The total number of offspring produced by the first male until the female ceased offspring production was 329 (27%) and by the second male was 894 (73%).

In the NS and SN groups, offspring paternity can be determined using the visible genetic markers carried by the swollen antenna of the Rd males. Overall, the pattern of P_2_ between a sibling and non-sibling male when competing against a Rd male was similar (estimate: -0.21, z = -0.48, P = 0.62, Fig. 3 panel B).

### Calculating adjusted P_**2**_

In those females with offspring from both males a *t*-test was used to compare the proportion of adult offspring sired by sibling males when they were the last male to mate with the proportion sired by non-sibling males in the same role. There was a significant difference between the two groups (*t* = 2.85, df = 34, *P* = 0.007 using arcsine transformed data) with non-sibling males siring a greater proportion of offspring than sibling males. However, because the offspring of eggs fertilised by sibling males are less likely to reach adulthood (as shown in the comparison of the number of offspring from NN and SS females in Figure 1), counting numbers of offspring will underestimate fertilisation success of sibling males. We adjusted P_2_ values to allow for this by dividing the number of adult offspring from the NN group by the number of adult offspring from the SS group (the answer being 1.42). This figure was multiplied by the number of offspring produced by each female from the sibling male. The adjusted numbers of adult offspring from sibling males (when second or first male to mate) were used to calculate an adjusted P_2_, which was 0.78 ± 0.04 for sibling males and 0.85 ± 0.027 for non-sibling males, a non-significant difference (*t* = 1.26, df = 34, *P* = 0.22 using arcsine transformed data).

## Discussion

Males were very active in soliciting copulations in both sex ratio treatments. As previously observed by [23], the vast majority of mating attempts were initiated by males, which attempted to mate as frequently with females (siblings and non-siblings) as with other males, suggesting that they cannot distinguish between the sexes. The similar number of homosexual and heterosexual copulations is what we would expect if only males initiate copulations; if males and females were equally active in initiating copulation attempts then we would expect twice as many heterosexual copulations as homosexual copulations in the double male treatment since there are 2 males and 1 female in each trial. The number of attempted matings and the proportion of attempts that were successful was higher when there was only one male present, suggesting that it is availability of females that limits attempted matings rather than male activity.

Males placed with a sibling and a non-sibling female attempted to mate with their sisters and actually succeeded in mating with them more often than unrelated females. This suggests that males can recognize their sisters. Given that there does appear to be substantial inbreeding depression in our population (Figure 1), and in the species in general [20], these findings are unexpected. Siblings in this experiment were separated as pupae, but had spent all their larval stages (about 3 weeks since hatching) together and hence had some experience of each other. It is also possible that they could identify siblings using the products of highly polymorphic genes [7]; we cannot distinguish between learned familiarity and genetic similarity. There are inclusive fitness benefits of mating with closely related individuals [42, 43], which mean that it is actually quite surprising that we do not see more examples of individuals preferentially mating with close relatives in nature [44]. One possibility is that *Tribolium* beetles with their relatively low costs of inbreeding are adaptively choosing to mate with relatives.

In relation to potential post-mating inbreeding avoidance, Females mated to 2 non-sibling males produced significantly more offspring than those mated to 2 sibling males or to a non-sib and then a sib male. This difference is as expected given the high P_2_ in this species [45, 46]; females have more viable offspring if they mate last to a sibling, but their offspring will not suffer very much if an earlier mate is their sibling. It appears that female *T. castaneum* do not bias paternity in favour of unrelated males, which is consistent with de Boer et al.’s [47] meta-analysis which found little support for inbreeding avoidance in animals in general. However, our study suggests that Tribolium could avoid costs of inbreeding if they could somehow ensure that the last male they mate with before laying eggs is an unrelated male. Similarly, a study on this species showed that, polyandry rescues female reproductive fitness within inbred populations [22]. As well as confirming the previously reported high P_2_ immediately following a double mating, our results show that P_2_ decreases significantly over a 5 week period. This is consistent with the suggestion by Lewis and Jutkiewicz [46] that high P_2_ is a result of sperm being stratified in the female spermatheca, but suggests that this stratification breaks down over time. Lewis and Jutkiewicz [46] used dissection of mated females to show that a single insemination only fills about two-thirds of the total capacity of the spermatheca, with the remaining one third being filled by the next copulation. Our results show that sperm displacement occurs during the second copulation otherwise we would expect that when a female uses all the sperm she has available, approximately two thirds of her offspring will be sired by the 1^st^ male to mate. We show that the majority of offspring produced by doubly mated females that ceased offspring production over the 5 weeks (presumably because they run out of sperm) were sired by the second male. This suggests that sperm displacement is a factor driving the high P_2_ seen in this species, a proposal which is consistent with Haubruge et al.’s [23] suggestion that the morphological characters of the male genitalia of *T. castaneum* may allow removal of previously deposited sperm from the female reproductive tract.

Although inbreeding avoidance mechanisms are taxonomically widespread and have been shown to be the rule rather than the exception amongst vertebrates [3], examples of either pre- or post-mating inbreeding avoidance in insects are rare. Field crickets *Gryllus bimaculatus* have been shown to prefer unrelated males as mates [48, 49] and appear to be able to avoid using sperm from closely related males [15, 16], and there is evidence that another species of cricket may show similar abilities [14]. The ant *Iridomyrmex humilis* avoids mating with kin [50], as do females of the predaceous mite *Phytoseiulus persimilis*, [51]. However the Argentine ant, *Linepithema humile* shows similar among and between sibling mating rates [52]. Our population of *T. castaneum* also failed to show pre- or post-mating inbreeding avoidance despite survival to adulthood being substantially reduced in inbred offspring and the potential for other adult fitness costs that we have not measured (e.g., Vasudeva et al. 2025). This situation has three potential explanations; either that inbreeding risk is not sufficient to drive the evolution of inbreeding avoidance, that inclusive fitness benefits of inbreeding outweigh the costs [41, 42] (as discussed above), or that there are constraints on the evolution of kin recognition. The risks of, and costs imposed by inbreeding depression in wild invertebrate populations remain important open questions, particularly in the light of evidence that costs of inbreeding may be amplified by thermal stress [53, 54].

